# Long-read *de novo* assembly of the red-legged partridge (*Alectoris rufa*) genome

**DOI:** 10.1101/2024.01.23.576805

**Authors:** Rayner González-Prendes, Ramona Natacha Pena, Cristobal Richart, Jesús Nadal, Roger Ros-Freixedes

## Abstract

The red-legged partridge (*Alectoris rufa*) is a popular game bird species that is in decline in several regions of southwestern Europe. The introduction of farm-reared individuals of a distinct genetic make-up in hunting reserves can result in genetic swamping of wild populations. Here we present a *de novo* genome assembly for the red-legged partridge based on long-read sequencing technology. The assembled genome size is 1.14 Gb, with scaffold N50 of 37.6 Mb and contig N50 of 29.5 Mb. Our genome is highly contiguous and contains 97.06% of complete avian core genes. Overall, quality of this genome assembly is equivalent to those available for other close relatives such as the Japanese quail or the chicken. This genome assembly will contribute to the understanding of genetic dynamics of wild populations of red-legged partridges with releases of farm-reared reinforcements and to appropriate management decisions of such populations.

## Background & Summary

The red-legged partridge (*Alectoris rufa*) is a popular game bird species from southwestern Europe (France, Spain, Portugal, and northwestern Italy). It can also be found in flat areas of England and Wales, where it was introduced as a game species. This partridge species can survive in a wide range of habitats, favouring dry low grass areas. Wild populations in the Iberian Peninsula are currently in decline^1,2^, mainly due to habitat degradation and hunting pressure. To meet the demand of birds for hunting purposes, farm-reared partridges are often released in game reserves.

Red-legged partridges belong to the Phasianidae family of the order Galliformes. Red-legged partridges and chickens (*Gallus gallus*) diverged about 65 million years ago^3^. Despite the relatively long divergence time, the genome synteny and karyotype of both species are highly conserved. The red-legged partridge has 2*n* = 78 chromosomes^3^ and the chicken has 2*n*= 80 in its latest genome assembly (GenBank accession: GCA_016699485.1). Most of the chromosomes are small microchromosomes, while only a few macrochromosomes are present in the karyotype^3–5^. This conservation in genome structure and organization between the red-legged partridge and the chicken, particularly on the macrochromosomes, is expected to facilitate the identification and characterization of orthologous genes and genomic regions.

The first red-legged partridge genome assembly^6^, published in 2021, was among the first to be done almost exclusively based on short-read sequencing data, and by current standards it could be considered of draft level with an N50 equal to 11.5 Mb^6^. The assembly included linkage data based on a low-density genetic map and the placement of scaffolds to chromosomes relied considerably on conserved synteny assumptions with the better assembled reference genome of the chicken. However, avian microchromosomes have proved to be difficult to assemble even with current technology.

An improved genome assembly is crucial for better understanding genomic variation in wildlife populations and commercial stocks. Improved annotation of coding and non-coding genes benefits the functional interpretation of genome-wide scans for selection sweeps and genome-wide association studies (GWAS), as well as aiding in the identification of genetic markers for assisting in selection of commercial stocks or gene editing targets. In wild populations, an improved genome assembly can also help in capturing population-specific variation that could be used in the design of conservation programs. So far, in the Galliformes, the genome assemblies of the chicken^7,8^, Japanese quail (*Coturnix japonica*)^9,10^, Gunnison sage-grouse^11^, helmeted guineafowl^12^, and turkey^13^ are high-quality genome assemblies based on long-read sequencing technology. These high-quality assemblies allow for comparative studies within the Galliformes and raise the opportunities for further research. In this study, we aimed to develop a new *de novo* assembly for the red-legged partridge species based on a combination of long- and short-read sequencing technologies. This new assembly will provide an improved reference genome that will be a useful resource for future research and practical applications, including conservation efforts.

## Methods

### Ethics statement

Sample collection was conducted in full compliance with Spanish laws and regulations, including the licence of “Las Ensanchas” for sampling shot partridges. The protocol was approved by the Committee on the Ethics of Animal Experiments of the University of Lleida (Ref. 1998-2012/05).

### Sample collection and construction of sequencing libraries

Muscle samples for whole-genome sequencing were collected from 60 red-legged partridges, divided in two subpopulations. Half of the samples (n=30) were from a population of wild partridges from a private game hunting state in central Spain (“Las Ensanchas”, Ciudad Real, Spain) and the other half (n=30) were from a partridge rearing farm (more information in Ros-Freixedes et al.^14^). These 60 samples were sequenced in an Illumina (San Diego, CA, USA) short-read platform, as described below. In addition, blood samples from two additional red-legged partridges, one from each subpopulation, were used for long-read sequencing in an Oxford Nanopore Technologies (Oxford, UK) platform. Total genomic DNA was isolated using standard protocols consisting of proteinase K lysis and phenol-chloroform purification^15^. The DNA for the long-read sequencing experiment was isolated using wide-bore filter tips and the preparations were never frozen before sequencing. The purified DNA was quantified using a Qubit fluorometer (Invitrogen, Waltham, MA USA) and quality checked using a NanoDrop Spectrophotometer (ThermoScientific, Wilmington, DE, USA) and agarose gel electrophoresis.

The short-insert paired-end libraries for the whole-genome sequencing were prepared with a PCR-free protocol using KAPA HyperPrep kit as detailed before^14^. The 60 libraries were sequenced on NovaSeq6000 (Illumina) in paired-end mode with a read length of 2x151+17+8 bp following the manufacturer’s protocol for dual indexing. Image analysis, base calling and quality scoring of the run were processed using the manufacturer’s software Real Time Analysis (RTA 3.4.4, Illumina) and followed by generation of FASTQ sequence files. For each sample, a minimum of 20 Gb of sequencing data was generated, which represents a sequencing depth of ∼20x of the partridge genome.

For the long-read sequencing experiment, one sequencing library was prepared from a pool of equal amounts of the two DNA samples. The library was built using the Ligation sequencing kit SQK-LSK109 from Oxford Nanopore Technologies. Briefly, 4.0 μg of the pool DNA was repaired and end-repaired using NEBNext FFPE DNA Repair Mix (New England BioLabs, Ipswich, MA, USA) and the NEBNext UltraII End Repair/dA-Tailing Module (New England BioLabs) and followed by the sequencing adaptors ligation, purified by 0.4X AMPure XP Beads and eluted in Elution Buffer (SQK-LSK109). The sequencing run was performed on GridION Mk1 (Oxford Nanopore Technologies) using a Flowcell R9.4.1 FLO-MIN106D (Oxford Nanopore Technologies) and the sequencing data was collected for 90 h. The quality parameters of the sequencing run were monitored by the MinKNOW platform version 3.6.5 in real time and basecalled with Guppy version 3.2.10.

### Genome assembly

For the genome *de novo* assembly we used the long-read sequencing data of the DNA pool as well as short-read sequencing data of one individual from the farm-reared subpopulation. We used Flye^16^ for initial assembly and NextPolish^17^ for the polishing step. This approach allowed us to combine the benefits of both long-read and short-read sequencing technologies to produce a high-quality assembly of the red-legged partridge genome. LRScaf^18^ software was then utilized to identify and close gaps in the assembly. The completeness of the final assembly in terms of gene space was evaluated using BUSCO^19^ v4.1.2, which was run in genome mode (-m genome) with the vertebrae (vertebrata_odb10) and avian (aves_odb10) datasets. The final draft assembly was aligned against the quail and chicken reference genomes using the default parameters in D-genie (https://dgenies.toulouse.inra.fr) using the Minimap2^20,21^ v2.26 aligner.

### Repeats annotation

We generated a *de novo* repeat library using the BuildDatabase tool from RepeatModeler^22^ v1.0.11. RepeatMasker^23^ v4.0.7 and a custom-built repeat library from RepeatModeler were used to comprehensively identify and annotate the repetitive elements in the genome. The custom RepeatModeler library and custom R scripts were used to investigate the differences in repeat content between the scaffolds. Each scaffold was split into bins (each bin corresponding to 2% of the scaffold length), allowing us to compare between the scaffolds by relative length.

### Gene prediction and annotation

The genome is currently being annotated using the ENSEMBL annotation pipeline, which is available as part of the Ensembl Rapid Release^24^.

### Alignment metrics

We assessed the quality of the alignment using the short reads from the 59 sequenced red-legged partridges (excluding the one used for genome *de novo* assembly) to our reference genome. To do so, DNA sequence short reads were pre-processed using Trimmomatic^25^ to remove adapter sequences from the reads. Then, we mapped the reads to our reference genome for *A. rufa* (Arufa2) as well as other already available reference genomes. The other reference genomes tested were a previously published reference genome for *A. rufa* (Arufa1^6^; GenBank accession: GCA_019345075.1) and those of two close poultry species: the Japanese quail (*C. japonica*; version 2.1; GenBank accession: GCA_001577835.2) and the chicken (*G. gallus*; version GRCg6a; GenBank accession: GCA_000002315.5). Additionally, we used data from 28 sequenced chicken mapped to the chicken reference genome as a control. To map the reads we used the BWA-MEM^26^ algorithm. Duplicates were marked with Picard (http://broadinstitute.github.io/picard). Alignment metrics were extracted using Picard.

### Genetic variation and inbreeding

In a previous study we used the reference genome of the quail to align the whole-genome sequence data of wild and farm-reared red-legged partridges to study their genetic variation, inbreeding, population structure and selective sweeps^14^. We benchmarked our new Arufa2 reference genome by comparing to Arufa1 and the quail reference genome. To do so, single nucleotide polymorphisms (SNPs) and short insertions and deletions (indels) were identified with the variant caller GATK HaplotypeCaller^27,28^ (GATK 3.8.0) using default settings with all three reference genomes. Variant discovery with GATK HaplotypeCaller was performed separately for each individual and then a variant set for the whole population was obtained by jointly genotyping all individuals at the variant sites with GATK GenotypeGVCFs. We retained all biallelic variants for further analyses with VCFtools^29^. To enable comparisons, only data from scaffolds that aligned to chromosomes 1 to 28 of the quail reference genome were considered for Arufa2 and Arufa1.

For the analysis of the transitions-to-transversions ratio (Ti/Tv), nucleotide diversity (π) and Tajima’s D^30^ we used all called variants to avoid biases caused by the alteration of the allele frequency spectrum. These parameters were calculated with VCFtools using non-overlapping windows of 10 kb for π and Tajima’s D. We calculated the fixation index (F_ST_) between both populations using Weir and Cockerham’s weighted estimator^31^ in the same 10-kb windows.

For the estimation of inbreeding coefficients and the detection of runs of homozygosity (ROH) we only used variants with minor allele frequency (MAF) ≥ 0.05 across both populations. We estimated the inbreeding coefficient (F_IS_)^32^ based on the homozygosity rate of each individual, using the method-of-moments implemented in the --het function in PLINK^33^ 1.9. We detected ROH using the --homozyg function in PLINK 1.9 with the parameters to define the ROH segments as proposed by Ceballos et al.^34^ for whole-genome sequence data; that is: --homozyg-snp 50 --homozyg-kb 300 --homozyg-density 50 -- homozyg-gap 1000 --homozyg-window-snp 50 --homozyg-window-het 5 --homozyg-window-missing 5 --homozyg-window-threshold 0.05. Inbreeding based on the ROH (F_ROH_) was calculated as the sum of the lengths of ROH relative to the total length of the chromosomes or scaffolds used.

## Data Records

The short-read sequencing data were submitted to the NCBI Sequence Read Archive (SRA) database with accession number SRP408849 and Bioproject accession PRJNA824288. The long-read sequencing data are available via ENA (Bioproject accession PRJEB67643, Biosample: ERS16499794, ERS16499793, ERS16499792, ERS16499791, ERS16499790).

The assembled draft genome of red-legged partridge has been deposited at GenBank under the accession number of GCA_947331505.1.

## Technical Validation

### Quality assessment of the genome assembly

The genome of the red-legged partridge was assembled using long-read data generated in a GridIon Nanopore platform and short-read data from the Illumina NovaSeq6000 sequencing system. Approximately three million long reads were produced with an N50 read length of 40,470 bp and a mean read length equal to 20,672 bp. Long reads were assembled and polished with short reads with the Flye^16^ assembler giving a 1.14 Gb genome draft size with 546 contigs (**Table 1**). Of the 546 contigs, 223 had a length equal or superior to 25 Mb and 168 to 50 Mb. The longest contig had a length of 111.5 Mb. The draft was scaffolded using LRScaf^18^ to produce a final assembly consisting of 426 scaffolds with a scaffold N50 of 37.6 Mb. The number of unassigned bases was 2,039.

**Table 1:** Assembly statistics of red-legged partridge genome using Nanopore long reads and Illumina short reads.

| Features | Arufa2 | Arufa1 | Cjaponica v2.1 | GRCg6a | GRCg7b* |
| --- | --- | --- | --- | --- | --- |
| Total sequence length (bp) | 1,142,509,937 | 1,027,480,606 | 927,656,957 | 1,065,348,650 | 1,053,332,251 |
| Total ungapped length (bp) | 1,141,765,707 | 996,170,856 | 917,263,224 | 1,055,564,190 | 1,049,948,333 |
| Gaps between scaffolds | 0 | 0 | 519 | 68 | 0 |
| No. of scaffolds | 426 | 10,598 | 2,531 | 524 | 214 |
| Scaffold N50 (bp) | 37,566,138 | 11,577,318 | 2,975,000 | 20,785,086 | 90,861,225 |
| Scaffold L50 | 9 | 25 | 86 | 12 | 4 |
| No. of contigs | 546 | 29,132 | 9,642 | 1,402 | 677 |
| Contig N50 (bp) | 29,494,169 | 97,167 | 511,217 | 17,655,422 | 18,834,961 |
| Contig L50 | 11 | 2,907 | 515 | 19 | 18 |
| Total number of chromosomes | - | - | 32 | 34 | 42 |
| GC (%) | 42 | 41 | 41 | 42 | 42 |
| Genome coverage depth (x) | 60 | 96 | 73 | 82 | 102 |

\* GRCg7b: *Gallus gallus*; GenBank accession: GCA\_016699485.1.

To evaluate the completeness and accuracy of the successive assembly versions, we used BUSCO^19^ and a whole-genome alignments approach. BUSCO results showed that all four assembly versions had over 96% of the expected vertebrate gene sets (**Fig. 1**). The four assembly versions that were tested included assembly produced with alternative software tools wtdbg2^35^ or Flye^16^, followed or not by a polishing step^17^. Among these assemblies, the assembly produced with Flye software and a polishing step emerged as the most comprehensive, boasting the highest number of complete BUSCOs coupled with a relatively low missing count (3,231 complete, 41 fragmented, and 82 missing BUSCOs).

**Figure 1:**
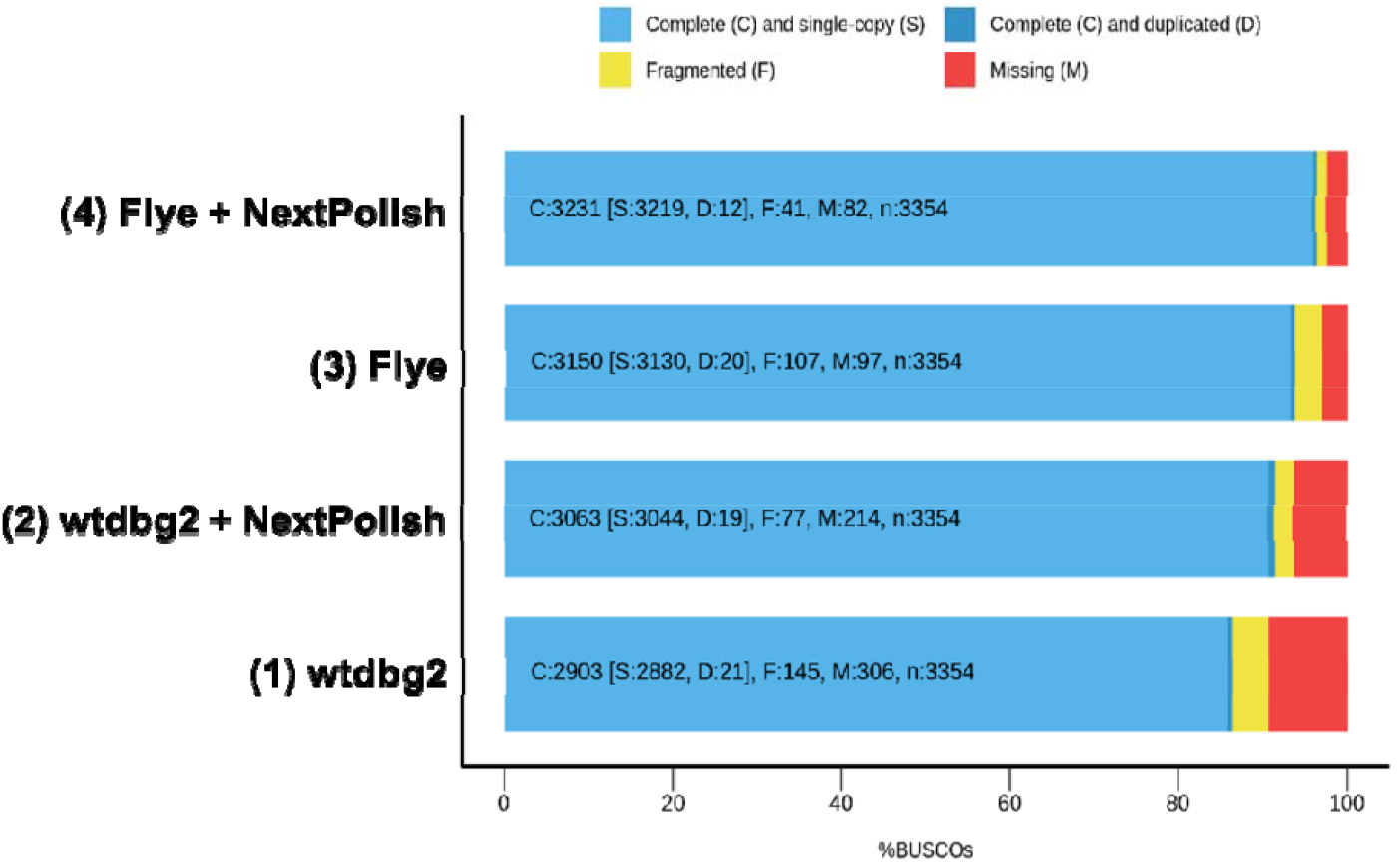
Assembly completeness measured in BUSCO scores. Percentage of aligned genes (x-axis) for the vertebrae (n=3,354) gene set in four red-legged partridge Arufa2 assembly versions. The percentages of complete (C) orthologs split between and single-copy (S) and duplicated (D) orthologs, fragmented (F) orthologs, and missing (M) orthologs are shown. Each bar (y-axis) represents a genome draft version for the Arufa2 genome: produced with alternative software tools wtdbg2, without (1) or with (2) a polishing step, and Flye, without (3) and with (4) a polishing step.

The proposed assembly, generated with Flye^16^ and polished, had the best BUSCO scores and, therefore, it was evaluated using the avian gene set. The completeness assessment results revealed that, out of 8,338 total avian core genes queried, 8,093 (97.06%) were detected as complete, while 8,134 (97.55%) were identified as either complete or partial. A total of 204 core genes were missing, representing 2.45% of the entire set. The average number of orthologs per core gene was 1.00, and only 0.38% of the detected core genes had more than one ortholog. These findings suggest that our genome is highly contiguous and, in terms of its expected gene content, near completion. Indeed, this assembly improved the quality of the first red-legged partridge genome assembly^6^, which identified 94.9% of single-copy avian orthologs (7,913 single-copy orthologs out of 8,338 proteins). The gene completeness assessment of our proposed assembly for *A. rufa* has higher values compared with prior findings including the chicken genome GRCg6a (95.4%^36^) and in the quail genome^10^, yet slightly inferior to the latest chicken genome GRCg7b, which boasts a completeness of 99.1%.

### Genome alignment comparison

The alignment comparing Arufa2 (**Fig. 2**) with the quail and chicken reference genomes reveals pronounced structural coherence. This is evidenced by a distinct correlation along the diagonal. Few regions exhibit misalignments, indicative of structural variations, inversions, or genomic rearrangements. Some isolated regions show denser clustering of alignments, hinting at areas of conservation or duplication. Sporadic off-diagonal alignments may suggest translocations or genomic modifications. Overall, most chromosomes or segments present consistent alignment, which underscores the strong genomic synteny between the genomes. In essence, there is substantial genomic similarity and structural coherence.

**Figure 2:**
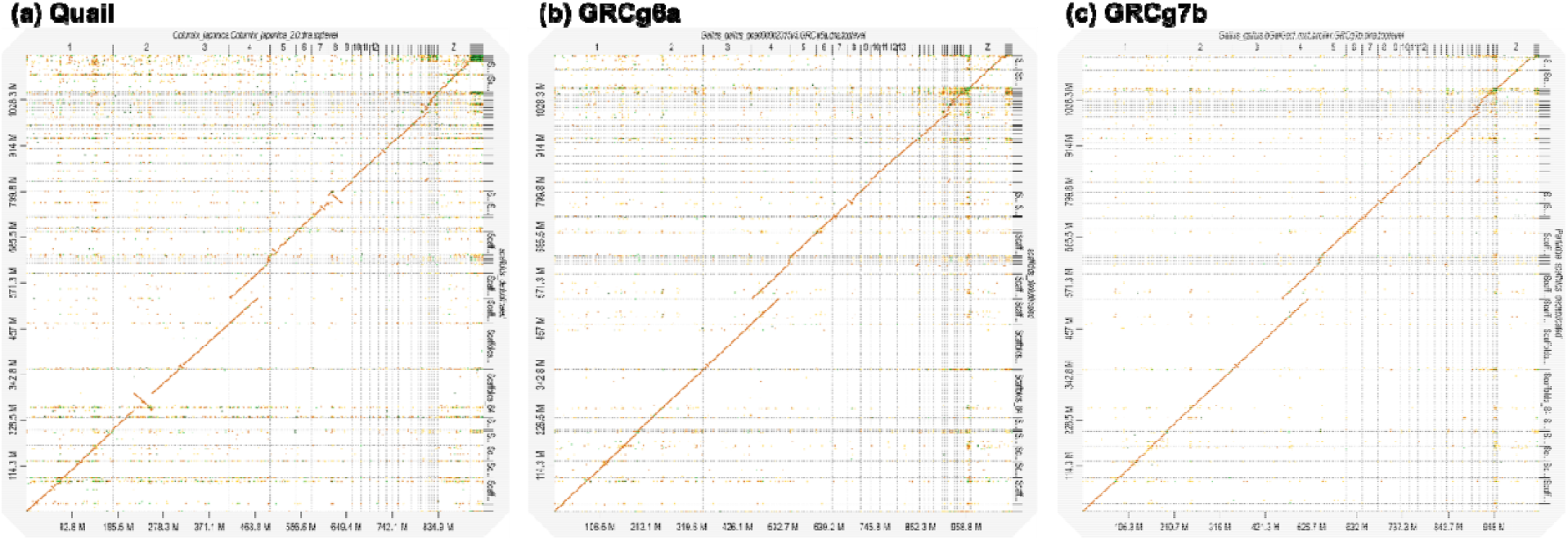
Genome alignment plots. Quail (a) and chicken, GRCg6a (b) and GRCg7b (c), genomes aligned with the red-legged partridge genome build. Alignment shows high structural coherence between genomes. The y-axis represents segments of our genome assembly in megabases (Mb) and the x-axis represents respective aligned segments in the counterpart genome. Orange dots indicate regions of alignment, with the diagonal line suggesting conserved genomic regions. Chromosomes are represented in the top horizontal bar, labelled from 1 to Z. These labels correspond to individual chromosomes of the *C. japonica* or *G. gallus* genomes. The side vertical bar provides a scale indicating the percentage of aligned scaffolds for each chromosomal segment.

To annotate repeats in the genome assembly, we used RepeatModeler^22^ to build a custom repeat library. Our analysis showed that repeats covered 10.45% of the genome, with the most common ones being LINE elements, which covered 6.35% of the genome. DNA transposons accounted for 0.76% of the bases, while LTRs and low complexity/simple repeats covered 0.53% and 1.58%, respectively. The remaining 1.23% of repeats were unclassified. These results provide insights into the repeat landscape of the red-legged partridge genome.

### Alignment metrics

Alignment metrics for the tested reference genomes are provided in **Table 2**. The value of long-read sequencing technologies is evident in the improved coherence of genome assemblies, although a larger proportion of reads with low mapping quality were detected in the scaffolded version of Arufa2 (13.9% excluded reads due to mapping quality scores) than in the non-scaffolded version (5.7%). Long-read sequencing presents a distinct advantage in structural variant detection. These reads span larger genomic regions, allowing for the accurate identification of insertions, deletions, inversions, and translocations that might be overlooked or inaccurately represented with short-read technologies^37,38^.

**Table 2:** Alignment metrics of 59 red-legged partridges sequenced with short reads against several reference genomes.

| Parameter* | Red-legged partridge reads mapped to: |  |  |  | Chicken reads mapped to |
| --- | --- | --- | --- | --- | --- |
|  | Arufa2 | Arufa1 | Cjaponica v2.1 | GRCg6a | GRCg6a |
| Before marking duplicates |  |  |  |  |  |
| Mean coverage depth (x) | 22.3 | 24.8 | 21.4 | 20.2 | 22.8 |
| SD coverage depth (x) | 67.0 | 139.4 | 91.5 | 69.3 | 63.1 |
| Het_Sensitivity | 0.984 | 0.996 | 0.909 | 0.877 | 0.975 |
| Het_Q | 18.2 | 23.8 | 10.4 | 9.0 | 16.0 |
| After marking duplicates |  |  |  |  |  |
| Mean coverage depth (x) | 17.7 | 21.7 | 18.3 | 17.6 | 19.4 |
| SD coverage depth (x) | 11.3 | 9.5 | 11.0 | 10.5 | 8.8 |
| Exc_MapQ (%) | 13.9 | 3.3 | 6.0 | 5.5 | 5.8 |
| Exc_Dupe (%) | 5.0 | 6.0 | 5.7 | 5.5 | 7.5 |
| Exc_BaseQ (%) | 1.3 | 1.4 | 1.3 | 1.3 | 1.4 |
| Exc_Overlap (%) | 14.2 | 15.9 | 15.0 | 15.0 | 13.8 |
| Exc_Capped (%) | 0.7 | 1.7 | 1.3 | 1.0 | 0.6 |
| Exc_Total (%) | 35.1 | 28.3 | 29.3 | 28.3 | 29.1 |
| Het_Sensitivity | 0.813 | 0.988 | 0.893 | 0.860 | 0.958 |
| Het_Q | 7.0 | 19.3 | 9.8 | 8.6 | 13.9 |

\* Mean coverage depth: mean coverage in bases of the genome territory, after all filters are applied. SD coverage depth: standard deviation of coverage of the genome after all filters are applied. Exc\_XXX: fraction of aligned bases that were filtered out because they were in reads with low mapping quality (score < 20; Exc\_MapQ), because they were in reads marked as duplicates (Exc\_Dupe), because they were of low base quality (score < 20; Exc\_BaseQ), because they were the second observation from an insert with overlapping reads (Exc\_Overlap), because they would have raised coverage above the capped value (250x; Exc\_Capped), or because all filters (Exc\_Total). Het\_Sensitivity: theoretical sensitivity for all heterozygous variants. Het\_Q: Phred-scaled Q score of the theoretical sensitivity for all heterozygous variants.

### Genetic variation and inbreeding

The Ti/Tv ratio was 2.50 with Arufa2, 2.54 with Arufa1, and 2.35 with Cjaponica. The Ti/Tv ratio was close to the generally expected values for whole-genome sequencing assuming spontaneous and neutral mutations^39^ and, in particular, almost identical to the value of 2.53 reported in chicken^40^. The number of biallelic called variants was 16,210,910, 17,659,943, and 69,609,618 with Arufa2, Arufa1, and Cjaponica, respectively. The SNP/indel ratio was 7.8 for both Arufa2 and Arufa1 and 5.8 for Cjaponica. However, most variants called with Cjaponica were very rare. A total of 55% of the variants called with Cjaponica were singletons (i.e., the minor allele was observed only once) compared to 27% with Arufa2 and Arufa1. The number (and SNP/indel ratio) of variants with MAF ≥ 0.05 was only of 6.494.820 (8.5), 7.037.619 (8.4), and 19,631,161 (4.8) with Arufa2, Arufa1, and Cjaponica, respectively. The use of a reference genome from a different species can result in an inflation of the number of called variants, as well as an enrichment of indels. These results are indicative of higher false positive rates when using a reference genome from a different species because indel calls are typically considered less reliable than SNP calls. In turn, this impacts the estimation of the other population genetic parameters.

The average nucleotide diversity (π) and Tajima’s D estimates within each population were very similar across all tested reference genomes (**Fig. 3a and 3b**). However, when we used the red-legged partridge reference genomes, we observed an enrichment of regions with low π estimates, which could be caused by selective sweeps that have removed variation in regions with low recombination rates. Thus, alignment to the quail reference genome may be less effective to detect such selective sweeps in red-legged partridge. Similarly, the average F_ST_ estimate was similar across reference genomes, but the quail reference genome increased the number of negative estimates (**Fig. 3c**). Negative Weir and Cockerham’s F_ST_ estimates may indicate greater variation within population than between populations, which is compatible with an inflation of false positive called variants.

**Figure 3:**
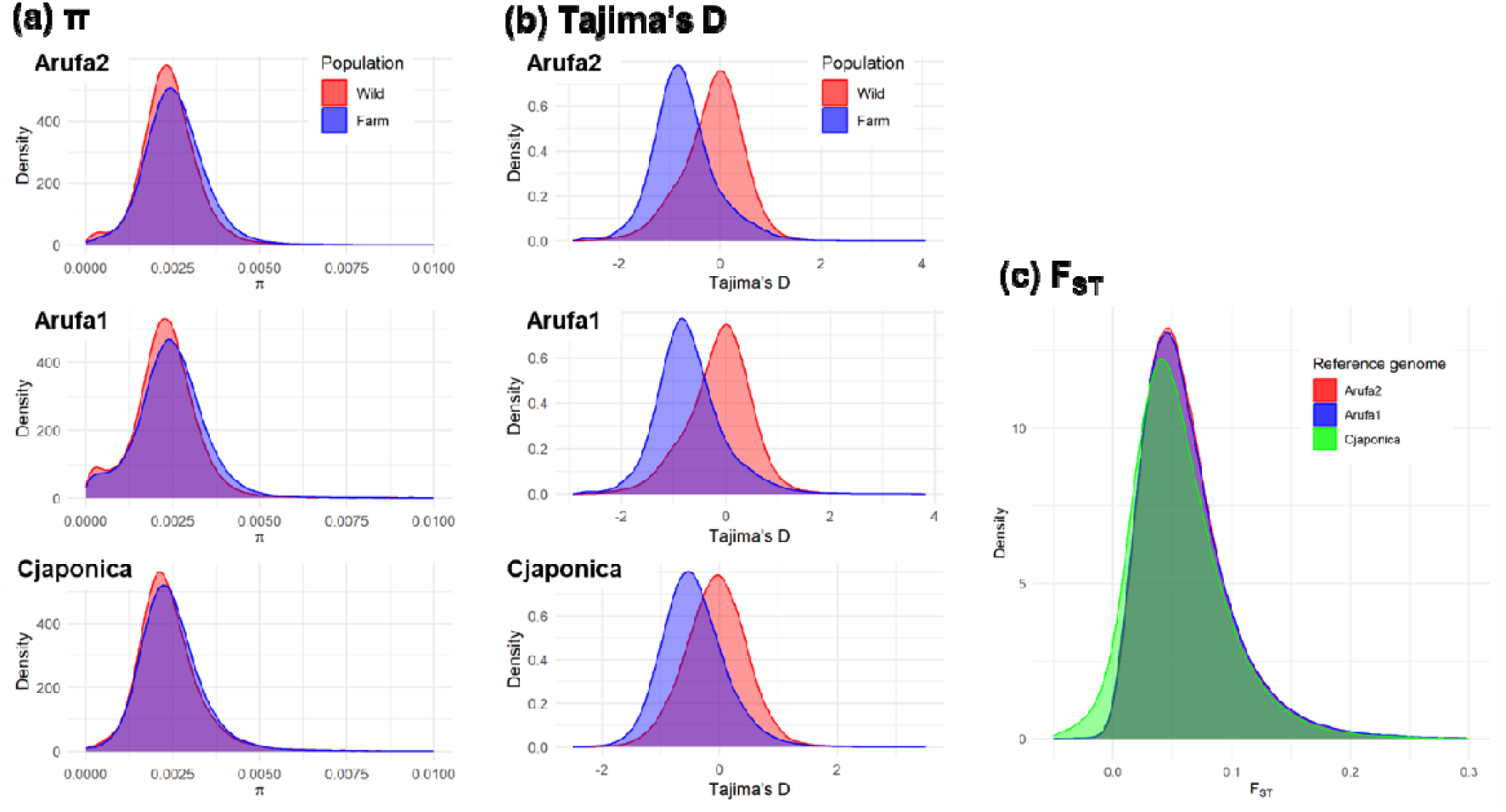
Distribution of nucleotide diversity (π), Tajima’s D, and fixation index (F_ST_) estimates in willd and farm-reared partridges with three different reference genomes.

Inbreeding coefficients based on the homozygosity rate were also likely overestimated when the quail reference genome was used compared to when a red-legged partridge reference genome was used (see F_IS_ in **Table 3**) because many of these variants will have a very low frequency of the alternative allele. At the same time, the greater high variant density and the large number of rare variants that arise from whole-genome sequencing hinder the detection of ROH when using a reference genome from a different species (see F_ROH_ in **Table 3**). Nonetheless, all inbreeding coefficients with all tested reference genomes support greater inbreeding for the farm-reared population than for the wild population, which is more consistent with expectations for captive-bred populations than our previous estimates using the quail reference genome^14^.

**Table 3:** Inbreeding coefficients in wild and farm-reared partridges based on homozygosity (F_IS_) or runs of homozygosity (F_ROH_) by reference genome.

| Inbreeding coefficients | Population |  | Welch's t-test |
| --- | --- | --- | --- |
|  | Wild | Farm-reared | p-value |
| Arufa2 |  |  |  |
| $F_{IS}$ | 0.026 (0.022) | 0.059 (0.027) | <0.001 |
| $F_{ROH}$ | 0.014 (0.011) | 0.020 (0.007) | 0.02 |
| Arufa1 |  |  |  |
| $F_{IS}$ | 0.022 (0.022) | 0.051 (0.027) | <0.001 |
| $F_{ROH}$ | 0.014 (0.012) | 0.019 (0.008) | 0.05 |
| Cjaponica v2.1 |  |  |  |
| $F_{IS}$ | 0.676 (0.007) | 0.687 (0.008) | <0.001 |
| $F_{ROH}$ | 0.000 (0.000) | 0.000 (0.000) | 0.08 |

Standard deviation (SD) is shown in parentheses.

Our findings underscore the importance of possessing a species-specific reference genome, such as that for the red-legged partridge. By using a dedicated reference, we can enhance the precision with which we estimate genetic population parameters. Such accurate estimations are instrumental in both characterizing and aiding conservation initiatives for wild populations of this species. Additionally, our reference genome presents a more refined scaffolding relative to previously available versions. Coupled with gene annotation, this opens the door to a range of other applications, notably the detection of selective sweeps.

## Code availability

The software versions and configurations employed are described below, following the steps of the pipeline available at https://github.com/CarolinaPB/nanopore-assembly with custom modifications:

1. Flye: --nano-raw raw.reads.fa.gz --out-dir draft_genome.fa --threads 16 and with wtdbg2 - x ont -g 4.6m -i raw.reads.fa.gz -t 16 -fo dbg
2. NextPolish: -genome draft_genome.fa -polish_type sr
3. LRScaf: -c draft_genome.fa -l long_reads.fq -o scaffolded_genome.fa
4. To retain a single copy of scaffolds and merge duplicated names, a Python script was utilized: ttps://github.com/CarolinaPB/Bioinfo_scripts/blob/main/remove_duplicates_fasta.py
5. BUSCO, version v4.1.2 was first run with the vertebrate database using the command: -m genome -l vertebrae (vertebrata_odb10) -i assembly.fa -o busco_output. It was then run again with the Avian orthologs gene set (aves_odb10).
6. RepeatModeler, version v1.0.11: BuildDatabase -name custom_repeat_db genome.fa
7. RepeatMasker, version v4.0.7: -lib custom_repeat_db genome.fa
8. ENSEMBL annotation pipeline: Uploaded the genome to ENSEMBL’s servers
9. The final draft assembly was employed as input for aligning with genomes from Figure 2 using the default parameters in D-genie (https://dgenies.toulouse.inra.fr) using the Minimap2 v2.26 aligner.

## Acknowledgements

We are grateful for the contributions made by the Melgarejo family, Patricia, Luis and Ivan Maldonado and Tom Gullick. Thanks also to the “Las Ensanchas” staff, especially the game keepers, the Barranquero family and collaborators, the members of the Tom Gullick hunting team in Campo de Montiel and around the world, Federación de Caza de Castilla y León, Delegación Burgalesa, MUTUASPORT, and Real Federación Española de Caza (RFEC). Carolina Ponz helped in sampling. Fundació Universitat Rovira i Virgili funded the sequencing (grant no. 2060-398-454-455; Proyecto IT20041-S; C. R.).

## Author contributions

**R. González-Prendes:** Methodology, Software, Validation, Investigation, Resources, Data curation, Writing – original draft, Writing – review and editing. **R**.**N. Pena:** Conceptualization, Methodology, Investigation, Resources, Writing – original draft, Writing – review and editing. **C. Richart:** Funding acquisition. **J. Nadal:** Conceptualization, Resources, Writing – review and editing, Funding acquisition. **R. Ros-Freixedes:** Conceptualization, Methodology, Software, Validation, Investigation, Resources, Data curation, Writing – original draft, Writing – review and editing.

